# Hat1-Dependent Lysine Acetylation Targets Diverse Cellular Functions

**DOI:** 10.1101/825539

**Authors:** Paula A. Agudelo Garcia, Prabakaran Nagarajan, Mark R. Parthun

**Author notes:** **Corresponding author:** Mark R. Parthun, PhD., 206 Rightmire Hall, 1060 Carmack Road, Columbus, OH 43210.

## Abstract

Lysine acetylation has emerged as one of the most important post-translational modifications, regulating different biological processes. However, its regulation by lysine acetyltransferases is still unclear in most cases. Hat1 is a lysine acetyltransferase originally identified based on its ability to acetylate histones. Using an unbiased proteomics approach, we have determined how loss of Hat1 affects the mammalian acetylome. Hat1^+/+^ and Hat1^−/−^ mouse embryonic fibroblast (MEF) cells lines were grown in both glucose- and galactose-containing media, as Hat1 is required for growth on galactose and Hat1^−/−^ cells exhibit defects in mitochondrial function. Following trypsin digestion of whole cell extracts, acetylated peptides were enriched by acetyllysine affinity purification and acetylated peptides were identified and analyzed by label-free quantitation. Comparison of the acetylome from Hat1^+/+^ cells grown on galactose and glucose demonstrated that there are large carbon source-dependent changes in the mammalian acetylome where the acetylation of enzymes involved in glycolysis was the most affected. Comparisons of the acetylomes from Hat1^+/+^ and Hat1^−/−^ cells identified 65 proteins whose acetylation decreased by at least 2.5-fold in cells lacking Hat1. In Hat1^−/−^ cells, acetylation of the auto regulatory loop of CBP was the most highly affected, decreasing by up to 20-fold. In addition to proteins involved in chromatin structure, Hat1-dependent acetylation was also found in a number of transcriptional regulators, including p53, and mitochondrial proteins. Hat1 mitochondrial localization suggests that it may be directly involved in the acetylation of mitochondrial proteins.

## INTRODUCTION

The acetylation of the ε-amino group of lysine residues in proteins is a widely utilized post-translational modification that regulates a variety of cellular processes, including chromatin structure, transcriptional regulation, DNA damage repair, cellular metabolism, cytoskeletal dynamics and apoptosis(1,2). The critical role played by protein acetylation is reflected in the number of human disease states, such as cancers and neurological disorders, which result from alterations in this modification(3). Proteomic studies have identified several thousand acetylated proteins in mammalian cells (known as the acetylome) (4–7). Given its abundance and relevance, acetylation may rival phosphorylation in importance as a regulatory and signaling modification. An intriguing difference between acetylation and phosphorylation regards the repertoire of enzymes responsible for these modifications. While there are more than 500 kinases encoded in mammalian genomes, there are fewer than 20 lysine acetyltransferases(1). Hence, it is likely that each known protein acetyltransferase will have multiple substrates and will be involved in multiple cellular processes(8,9).

Histone acetyltransferase 1 (Hat1) was the first protein ε-amino group lysine acetyltransferase to be identified and is the most widely conserved among eukaryotes (10–12). Like most lysine acetyltransferases, Hat1 was identified based on its ability to acetylate histones. Hat1 was originally purified from yeast cytoplasmic extracts in a complex with Hat2 (Rbap46/Rbbp7 in mammalian cells). Biochemically, the Hat1 complex acetylates histone H4 lysine residues 5 and 12 and only acetylates histones free in solution (the Hat1 reaction is blocked by the presence of nucleosomal DNA)(11,13). This observation suggested that Hat1 was responsible for the evolutionarily conserved diacetylation of newly synthesized histone H4 (on lysines 5 and 12) that is known to accompany the process of replication-coupled chromatin assembly, a hypothesis that was subsequently confirmed(14).

In vivo studies of Hat1 have identified several phenotypes associated with loss of this enzyme. These phenotypes, which include defects in heterochromatin gene silencing, DNA damage sensitivity, genome instability and alterations in gene expression, are likely due to the histone modifying function of Hat1(14–21). However, recent studies have identified mitochondrial defects associated with Hat1 loss, suggesting that this enzyme may influence key cellular pathways through the acetylation of non-histone substrates(22). To determine whether Hat1 influences non-histone protein acetylation, we have used a proteomic strategy to identify Hat1-dependent sites of acetylation. The abundance of acetylated peptides was compared between Hat1^+/+^ and Hat1^−/−^ mouse embryonic fibroblasts (MEFs) grown on both glucose and galactose. As previously reported in liver from mice fed a high fat diet or starved; these experiments indicated that the shift in carbon source from glucose to galactose had a dramatic effect on the acetylome(23,24). In particular, there were changes in the acetylation of many of the enzymes required for glycolysis. In addition, loss of Hat1 resulted in a significant decrease in the acetylation of >70 proteins. These proteins are involved in a wide range of cellular functions, including chromatin structure, transcriptional regulation and mitochondrial function. Interestingly, acetylation levels of the protein acetyltransferase CBP were the most highly affected in Hat1^−/−^ MEFs suggesting that Hat1 may be a regulator of CBP function. In addition, we demonstrate that Hat1 exhibits mitochondrial localization consistent with the possibility that this enzyme may be directly involved in mitochondrial protein acetylation.

## MATERIALS AND METHODS

### MEF cell culture

E14.5 embryos were dissected from uterus; head and internal organs were removed. Tissue was disaggregated using an 18-gauge syringe and brought to single cell suspension with trypsin incubation at 37°C. Cells were then plated onto 100 mm tissue culture plates, passaged upon confluency and maintained in Dulbecco’s modified Eagle medium (DMEM-Sigma) with 10% fetal bovine serum (FBS-Gibco) and 1X Pen/Strep antibiotics (Sigma). SV40 T immortalized MEFs (iMEFs) were derived from primary Hat1^+/+^ and Hat1^−/−^ embryonic day 13.5 embryos. To establish iMEFs, early passage cells were transformed with SV-40 T antigen containing plasmid pBSSVD2005 (ADDGENE, Cambridge, MA). Early passage cells were seeded at 25% confluency in 6 well plates and transfected with 2 ug of expression vector using Fugene reagent (Roche). Cells were harvested and seeded into 100 mm dishes after 48 hrs of transfection. The cells were split at 1 in 10 dilutions until passage 5.

### Mitochondrial Isolation

Cells were resuspended in mitochondrial isolation buffer (0.8M sucrose, 20mM tris-HCl ph7.2, 40mM KCl, 2mM EGTA, 1mg/mL BSA, 0.2mM PMSF), kept on ice for 2 min and lysed with Dounce homogenizer. Homogenate was centrifuged at 1500g for 10min at 4°C. Supernatant was further centrifuged at 17.000g for 30min at 4°C. Pellet was resuspended in mitochondrial extraction buffer without BSA.

### Subcellular Fractionation

Protein fractions from nucleus, cytosol and mitochondria were obtained according to Dimauro et al.(25). Cells were resuspended in STM buffer (250mM sucrose, 50mM Tris-HCl pH 7.5, 5mM MgCl2) and homogenized at low speed for 1 min, vortexed and kept on ice for 30min. Samples were then centrifuged at 800g for 15min, supernatant and pellet were separated; pellet was washed twice with STM buffer and resuspended in NET buffer (20mM Hepes pH7.9, 1.5mM MgCL2, 0.5NaCl, 0.2mM EDTA, 20% glycerol and 1%triton x-100) this was saved as the nuclear fraction. Supernatant was centrifuged one more time at 11.000g for 10min. Supernatant was treated with 100% acetone and centrifuged at 12.000g for 5min to obtain the cytosolic fraction. Pellet containing the mitochondrial fraction was washed twice with STM buffer and resuspended in SOL buffer (50mM tris-HCl pH 6.8, 1mM EDTA, 0.5% Triton X-100).

### Immunofluorescence

Hek293 cells were seeded on 6 well plates containing coverslips. Cells were fixed with 4% paraformaldehyde, blocked with 5% BSA and primary antibodies Hat1 (Abcam) was incubated overnight, washed and replaced with goat-anti-rabbit secondary antibody. Cells were mounted using Vectashield mounting media containing DAPI and imaged using a wide field fluorescence microscope. Mitolight was used according to manufacturer’s instructions.

### PTM scan lysine acetylation

2×10^8^ cells for each experimental condition were collected, the cell number came from pooling equal number of cells coming from three different wild type or Hat1 knock out cell lines. Cells were grown either on high glucose (25mM) DMEM or in glucose free DMEM supplemented with 10mM galactose for at least 4 days before collection. Cells were washed with PBS and resuspended in 10mL of PTMScan® urea lysis buffer (20 mM HEPES pH 8.0, 9.0 M urea, 1 mM sodium orthovanadate, 2.5 mM sodium pyrophosphate and 1 mM ß-glycerol-phosphate.). Cell lysates were frozen in dry ice/ethanol for at least 30 minutes, or until the cell lysate was completely frozen. The acetylome of each sample was analyzed by PTMScan (cell signaling) using their standard procedure. The log_2_ fold change was calculated using the relative fold change between the integrated peak area of the experimental conditions (Hat1^+/+^ galactose, Hat1^−/−^ glucose or Hat1^−/−^ galactose) and the control (Hat1^+/+^ glucose). Values greater than +/− 2.5 were used for subsequent analysis.

## RESULTS

Hat1 plays a role in multiple cellular processes through the acetylation of histone H4 and the regulation of chromatin structure(26–28). In addition, it has been reported that Hat1 is required for mitochondrial function(22,29). Given the large number of orphan acetylated proteins observed in mitochondria and the fact that Hat1 acetylates histone H4 in the cytosol, we decided to identify potential non-histone substrates of Hat1 and its additional functions. To do so, we took a proteomic approach to determine how loss of Hat1 influenced the mammalian acetylome.

### The mammalian acetylome is altered by a shift from glucose to galactose

Since it has been previously reported that metabolic stress can induce changes in acetylation and consistent with a role for Hat1 in mitochondrial function, we used two different growth conditions: glucose and galactose, to characterize the cellular acetylome from immortalized Hat1^+/+^ and Hat1^−/−^ MEFs(22–24).

Immortalized Hat1^+/+^ and Hat1^−/−^ MEFs were grown in media containing either 25mM glucose or 10mM galactose as the primary carbon source. Whole cell extracts were prepared, and proteins were digested with trypsin. Acetylated peptides were enriched by affinity purification using an anti-acetyl lysine affinity resin. The resulting peptides were analyzed by LC-MS/MS and the abundance of the acetylated peptides in each sample was determined using label free quantitation.

We first compared Hat1^+/+^ cells grown in glucose and galactose to determine the effect of the carbon source on the mammalian acetylome. 470 peptides, derived from 364 proteins were detected, whose level of acetylation changed by +/− 2.5-fold in cells grown in galactose relative to glucose (Figure 1A). Acetylation levels increased on 173 of these peptides (37%) and decreased on 297 (63%). Detailed information on all of the peptides identified in this study can be found in Supporting Table 1. Ingenuity Pathway Analysis was used to analyze the list of proteins whose acetylation changes by +/− 2.5-fold. Pathways showing a significant enrichment are listed in Figure 1B. Two of the significantly enriched pathways involve nuclear receptor signaling (TR/RXR activation and VDR/RXR activation). The primary proteins in these pathways that whose acetylation is altered by carbon source change are the protein acetyltransferase p300 and several nuclear receptor coactivators and repressors (Ncoa2, Ncoa3 and Ncor2).

**Figure 1.**
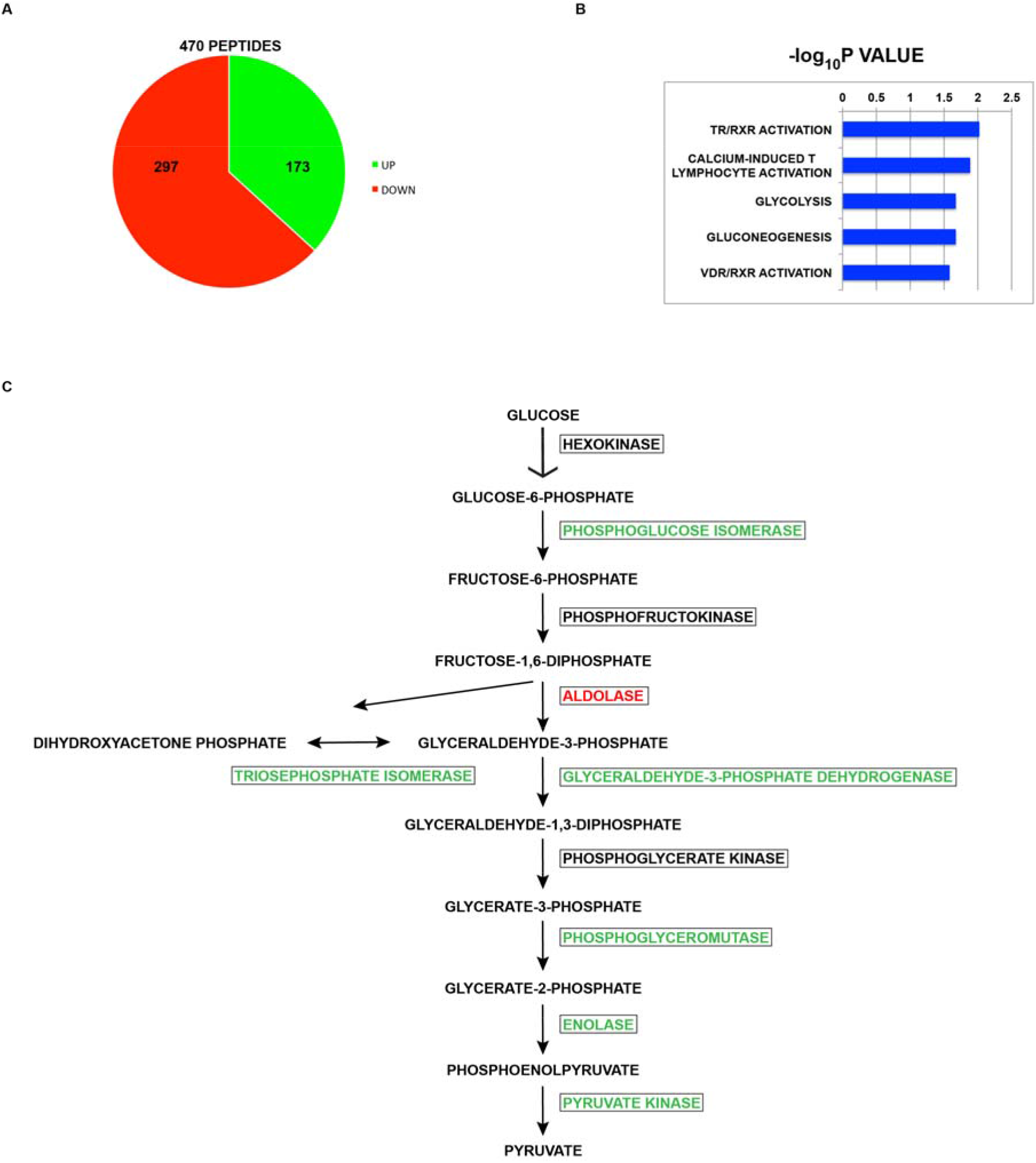
The effect of carbon source on the mouse acetylome. A) Pie chart showing the number of sites of lysine acetylation that increase or decrease when mouse embryonic fibroblasts are switch from growth in glucose-containing media to galactose-containing media. B) Ingenuity Pathway Analysis of the proteins whose acetylation changed by at least 2.5-fold between growth on glucose and galactose. The list of all peptides detected was used as the reference set. C) Schematic diagram of the pathway of glycolysis. Enzymes in green increased at least 2.5-fold when cells on galactose, enzymes in red decreased at least 2.5-fold on galactose and enzymes in black were unchanged.

Interesting, two other significantly enriched pathways are involved in glucose metabolism, glycolysis and gluconeogenesis (which share many of the same enzymes). We did not detect peptides from the enzymes needed to convert galactose to glucose-6-phosphate. Of the 10 enzymes required to convert glucose to pyruvate, 7 showed a significant change in acetylation. It has been shown that most of the enzymes involved in glycolysis, as well as enzymes in the majority of metabolic pathways, are acetylated(2,30). In fact, some of these sites of acetylation are known to regulate enzymatic activity. For example, acetylation of lysine 305 of pyruvate kinase promotes the enzyme’s degradation(31). However, none of the sites of acetylation that were found to change in response to growth on galactose have been the subject of functional characterization.

We also analyzed the acetylomes between Hat1^−/−^ MEFs grown in glucose and galactose. Table 1 lists the 50 proteins that had the largest increases and decreases in acetylation in Hat1^+/+^ MEFs and compares them to the changes observed in Hat1^−/−^ counterparts under the same growth conditions. It is apparent that loss of Hat1 had very little impact on the acetylation status of proteins whose acetylation decreased when grown on galactose. However, there were marked changes in the proteins whose acetylation increased in wild type cells grown in galactose. For 40% of these proteins, the increased acetylation seen when the cells were switched to galactose decreased by at least 5-fold in the Hat1^−/−^ cells. For example, PDZK1 showed a 108-fold increase in acetylation when Hat1^+/+^ MEFs were grown on galactose compared to only a 2.9-fold increase in Hat1^−/−^ MEFs. This set of proteins was not enriched for any specific cellular pathways but it includes 2 proteins that are critical for mitochondrial function, acylglycerol kinase (AGK) and ATP6V1E1. Interestingly, AGK^−/−^ cells have been reported to have very similar phenotypes to Hat1^−/−^ cells, including poor growth on galactose media, defective mitochondrial structure and poor mitochondrial metabolism(32). One possibility is that Hat1 mitochondrial function is executed through the acetylation of AGK regulating its activity.

**Table 1:**
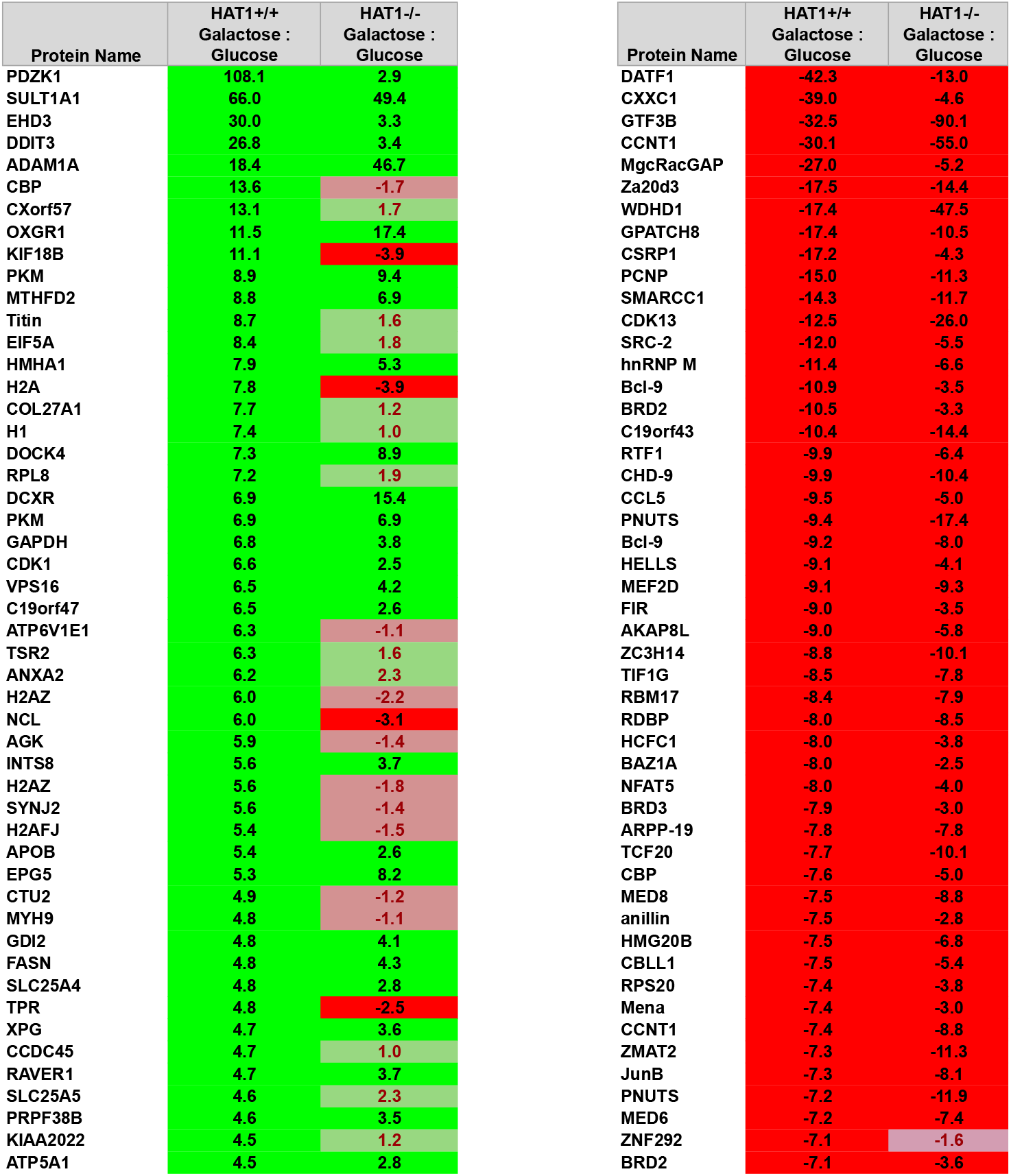
Top 50 proteins whose acetylation increases (green) or decreases (red) during growth on galactose relative to growth on glucose.

### Identification of Hat1-dependent sites of acetylation

Hat1-dependent sites of acetylation were identified by comparing the acetylome of Hat1^+/+^ MEFs grown on glucose to that of Hat1^−/−^ MEFs grown on glucose and the acetylome of Hat1^+/+^ MEFs grown on galactose to that of Hat1^−/−^ MEFs grown on galactose. This analysis identified 84 sites in 65 proteins whose acetylation decreased by at least 2.5-fold in Hat1^−/−^ cells (Table 2). Of these 84 sites of acetylation, 29 have not been previously reported (~35%, Table 2). Approximately 90% of these Hat1-dependent sites of acetylation were identified in cells grown on galactose media (Table 2). It was previously proposed that Hat1 acetylates lysine residues in the context of the sequence GXG**<u>K</u>**XG based on a comparison of the ability of Hat1 to acetylate histone H4 and H2A from a variety of eukaryotes(11). The sequence context of the Hat1-dependent sites of acetylation was analyzed to determine if there was an associated motif. Figure 2A shows the motifs identified for the total pool of acetylated peptides analyzed and for those peptides that changed by more than 2.5-fold in the Hat1^−/−^ samples. It is clear that glycine residues were enriched at the −3, −1 and +1 positions, as originally predicted. This motif also indicated that there was a strong preference for alanine at the +5 position and for glycine at the −7 position in the peptides whose acetylation was Hat1-dependent.

The list of proteins whose acetylation is Hat1-dependent was subjected to Gene Ontology (GO) analysis. Hat1-dependent protein acetylation was observed on proteins involved in a variety of cellular pathways, including ones not directly related to chromatin transactions (Figure 2B). These pathways included transcriptional regulation, translation, mitochondria and RNA processing.

**Table 2:**
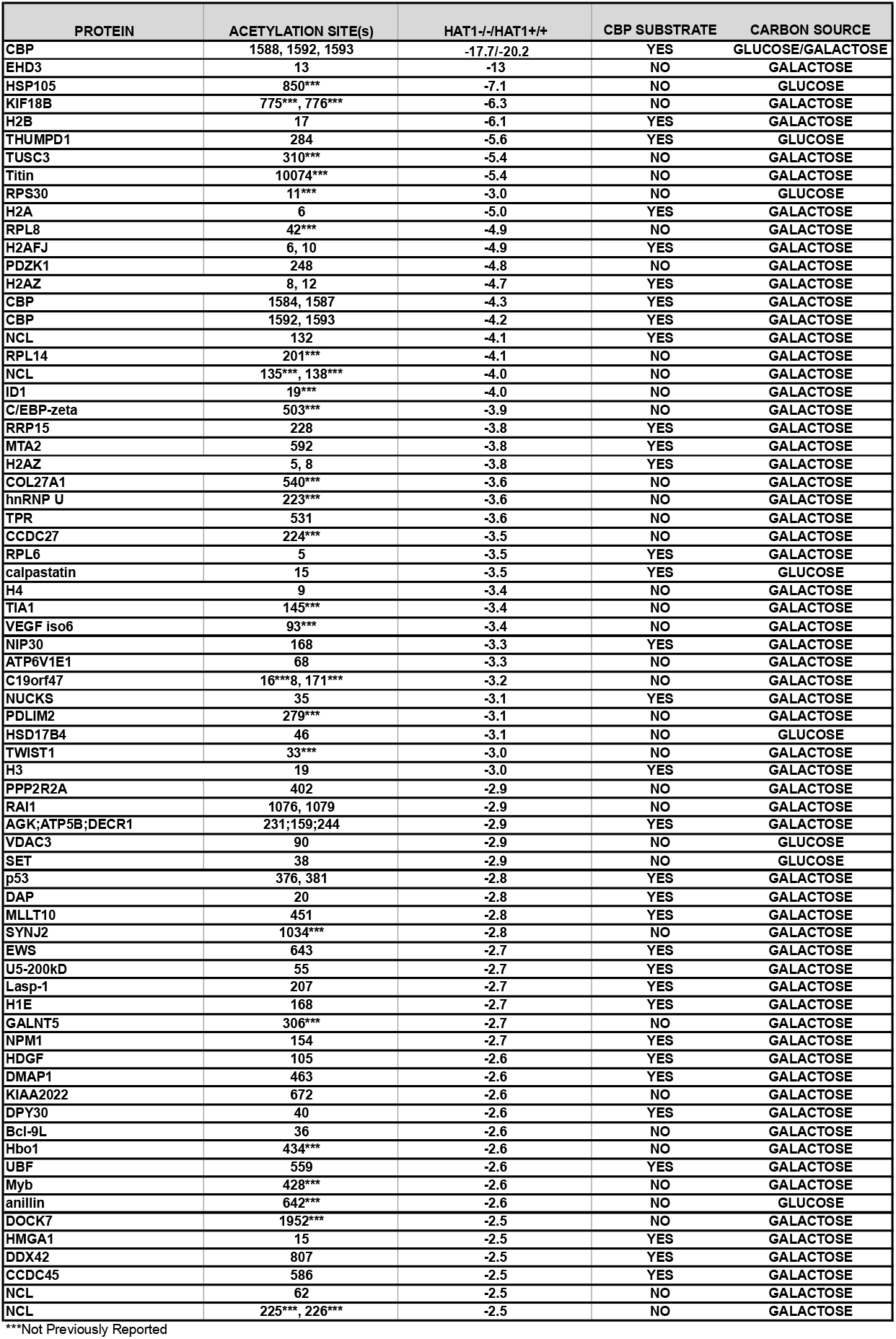
Acetylation sites that decrease by at least 2.5-forld in Hat1^−/−^ cells relative to Hat1^+/+^ cells.

**Figure 2.**
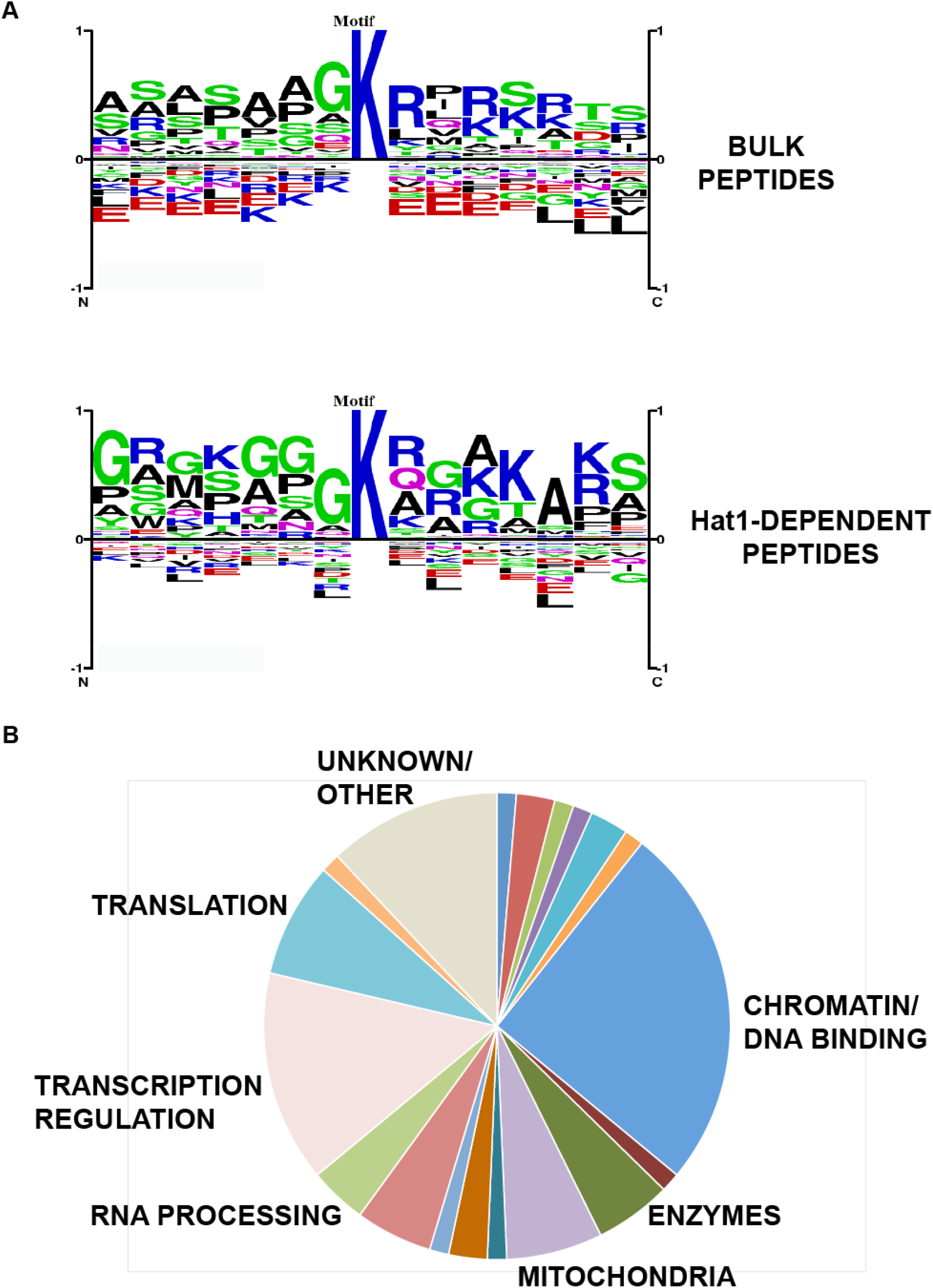
Characterization of the Hat1-dependent acetylome. A) Top, motif derived from the bulk acetylated peptides. Bottom, Motif derived from comparison of peptides whose acetylation is effected by Hat1 deletion. B) Proteins whose acetylation decreased at least 2.5-fold in Hat1^−/−^ cells were analyzed by Gene Ontology. The number of proteins involved in the indicated pathways are presented in a pie chart.

### Hat1 and the acetylation of chromatin related proteins

As expected, the most highly represented group of proteins whose acetylation was Hat1-dependent was those involved in chromatin and DNA binding (Figure 3A). Hat1-dependent acetylation was observed for all of the core histones, as well as for histone H1.4. These sites of acetylation included histone H2A lysine 5 (and the corresponding residues in histone H2A.Z and H2A.J) that was previously identified as a Hat1 substrate(11,33). Histone H4 lysine residues 5 and 12, the most well characterized Hat1 substrates, were not identified in these data sets. This is likely due to the abundance of trypsin cleavage sites in the NH_2_-terminal tail of H4 that results in the generation of fragments containing these residues that are too small to be reliably observed.

**Figure 3.**
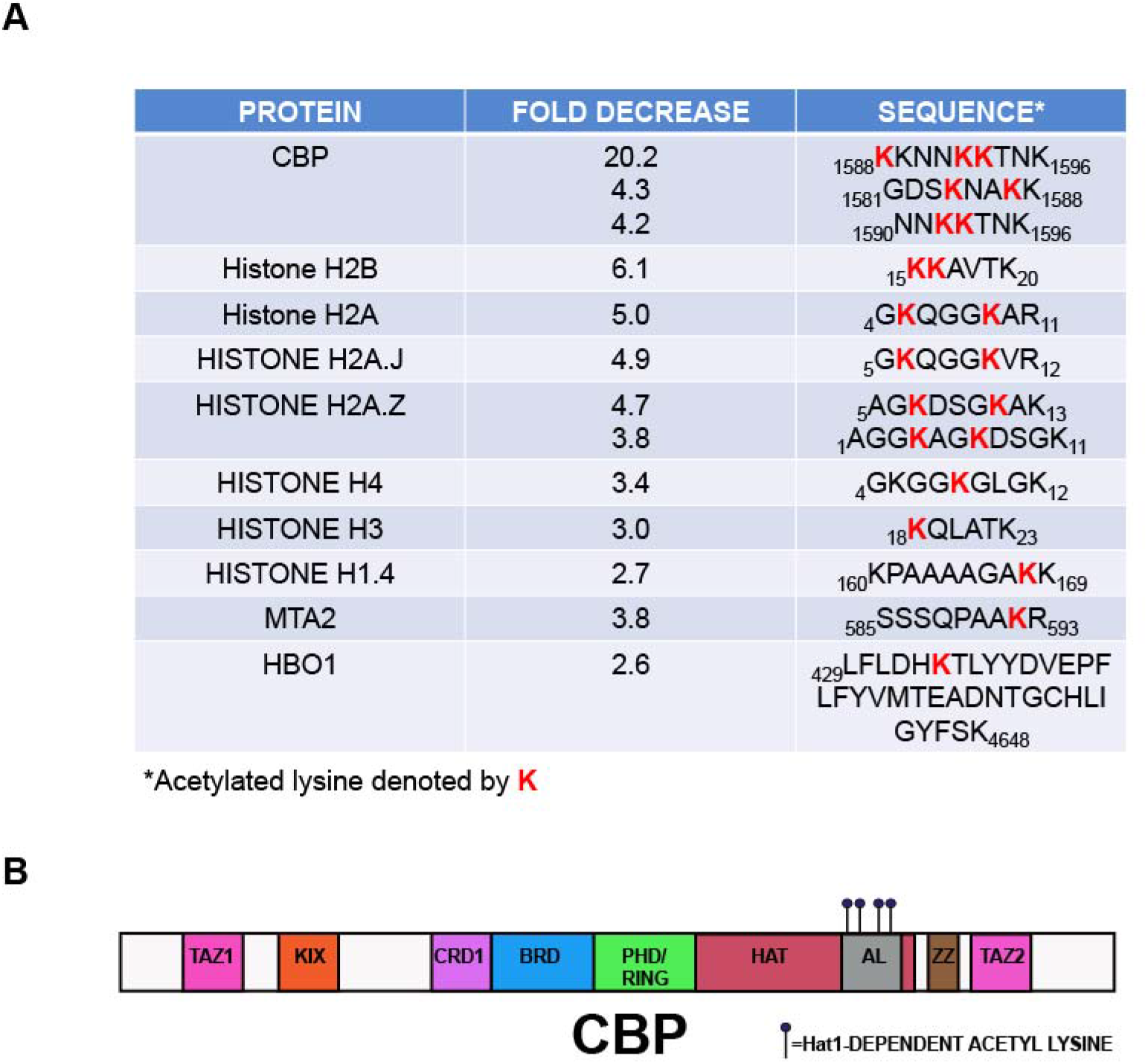
Hat1-dependent acetylation of chromatin proteins. A) A subset of chromatin-related proteins whose acetylation decreased at least 2.5-fold in Hat1^−/−^ cells. B) Schematic diagram showing the domains of CBP. Hat1-dependent sites of acetylation cluster in the autoregulatory loop.

Interestingly, the acetylation of two histone acetyltransferases (CBP and Hbo1) was Hat1-dependent (Table 2). The protein whose acetylation was most highly dependent on Hat1 was CBP, decreasing up to 20-fold in Hat1^−/−^ cells (Figure 3A). Importantly, the Hat1-dependent sites of acetylation on CBP are all located in the auto-regulatory loop (AL) in the CBP HAT domain (Figure 3B). The acetylation state of the AL of CBP is critical for its acetyltransferase activity(34). The dependence of the CBP AL on Hat1 suggests that Hat1 may influence the acetylation of some proteins indirectly through the activation of CBP. In fact, comparison of the Hat1-dependent sites of acetylation with an acetylomics analysis of CBP^−/−^ MEFs shows that approximately 45% of these sites are CBP substrates (Table 2)(8). The potential regulation of CBP by Hat1 is particularly intriguing given the role of CBP in the acetylation of newly synthesized histone H3(35). Several reports have suggested that Hat1 is involved in the transcriptional regulation of several genes in different models(36–40). However, this evidence is conflicted by Hat1’s incapacity to acetylate histones in a nucleosome context. One scenario is that Hat1 influences gene expression indirectly through the acetylation of CBP regulating the Hat activity of this transcription factor.

### Hat1 dependent protein acetylation of transcriptional regulators

Transcriptional regulatory proteins were also highly represented in the Hat1 acetylome (Figure 4A). An antibody specific for this site of p53 acetylation is available allowing us to validate the results of the proteomic analysis. We found that mouse p53 lysine 381 acetylation is induced by growth of MEFs in galactose (Supporting Table 1). Figure 4B shows immunofluorescence images of Hat1^+/+^ and Hat1^−/−^ MEFs grown in galactose. Staining for total p53 indicated that Hat1 loss did not affect p53 protein levels. However, there was a significant decrease of p53 lysine 381 acetylation in the Hat1^−/−^ MEFs, confirming that Hat1 regulates p53 acetylation.

**Figure 4..**
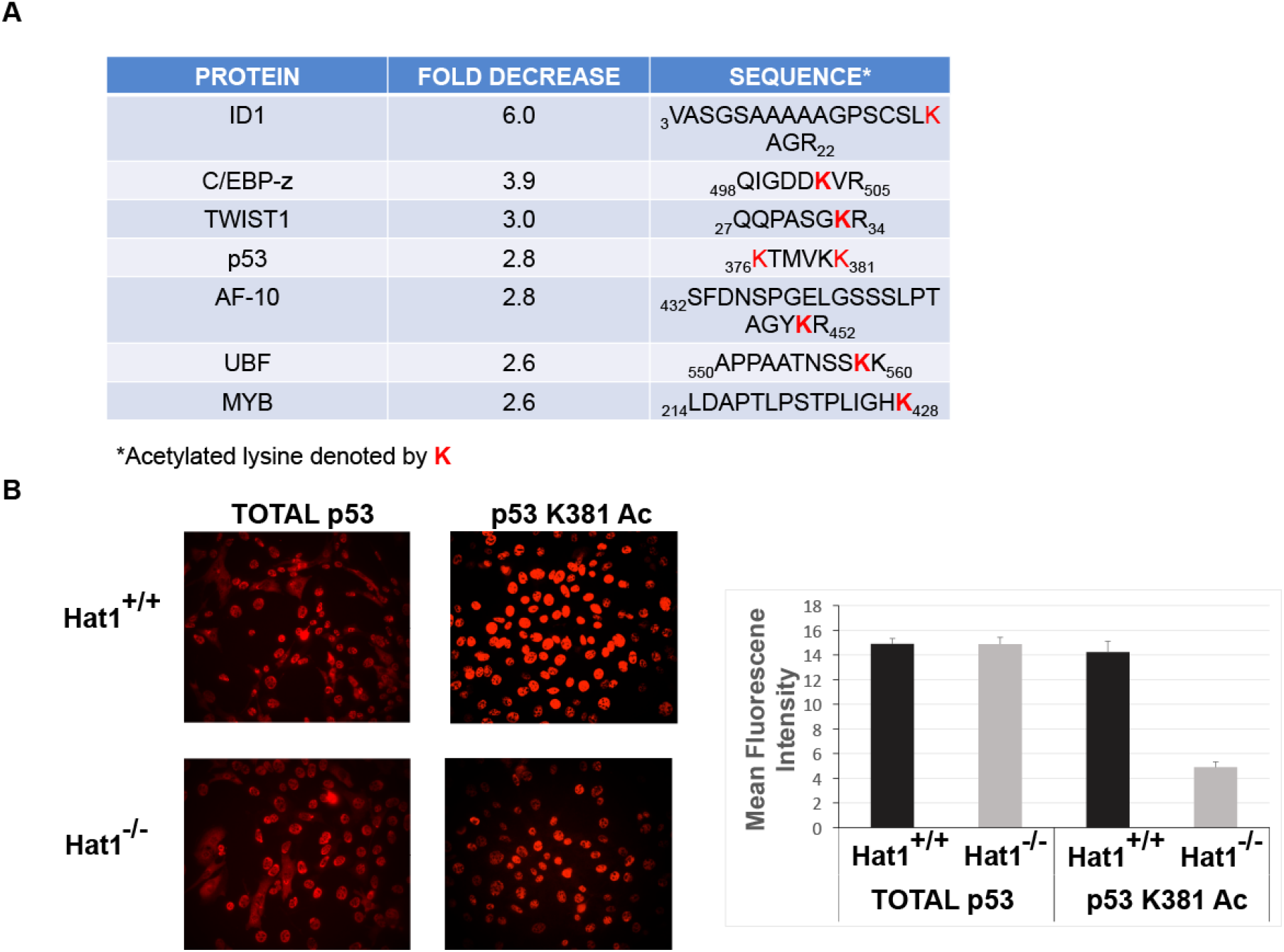
Hat1-dependent acetylation of transcriptional regulators. A) Subset of transcriptional regulators whose acetylation decreased at least 2.5 fold in Hat1^−/−^ cells. B) Hat1^+/+^ and Hat1^−/−^ were grown in galactose and stained with antibodies recognizing unmodified p53 and p53 acetylated at lysine 381 (386 in human). Fluorescent signals were quantitated (right).

### Hat1 and the acetylation of mitochondrial proteins

Proteomic approaches have identified a large number of mitochondrial proteins as being acetylated(5,41,42). Mitochondrial protein acetylation increases in mouse liver during fasting, caloric restriction or on a long-term high fat diet, indicating that the acetylation patterns in this organelle can respond to dietary changes(23,43–46). Interestingly, mitochondrial proteins are also enriched in the Hat1-dependent acetylome (Figure 5A). Remarkably, it is impossible to definitively assign one peptide as isobaric peptides are present in three distinct mitochondrial proteins; AGK, ATP5B and DECR1 (2,4-dienoyl CoA reductase). Recent reports have clearly identified defects in mitochondrial function in the absence of Hat1(22,29,47). The observation that the acetylation of a subset of mitochondrial proteins is Hat1-dependent suggests the possibility that Hat1 may influence mitochondrial function through mitochondrial protein acetylation.

**Figure 5.**
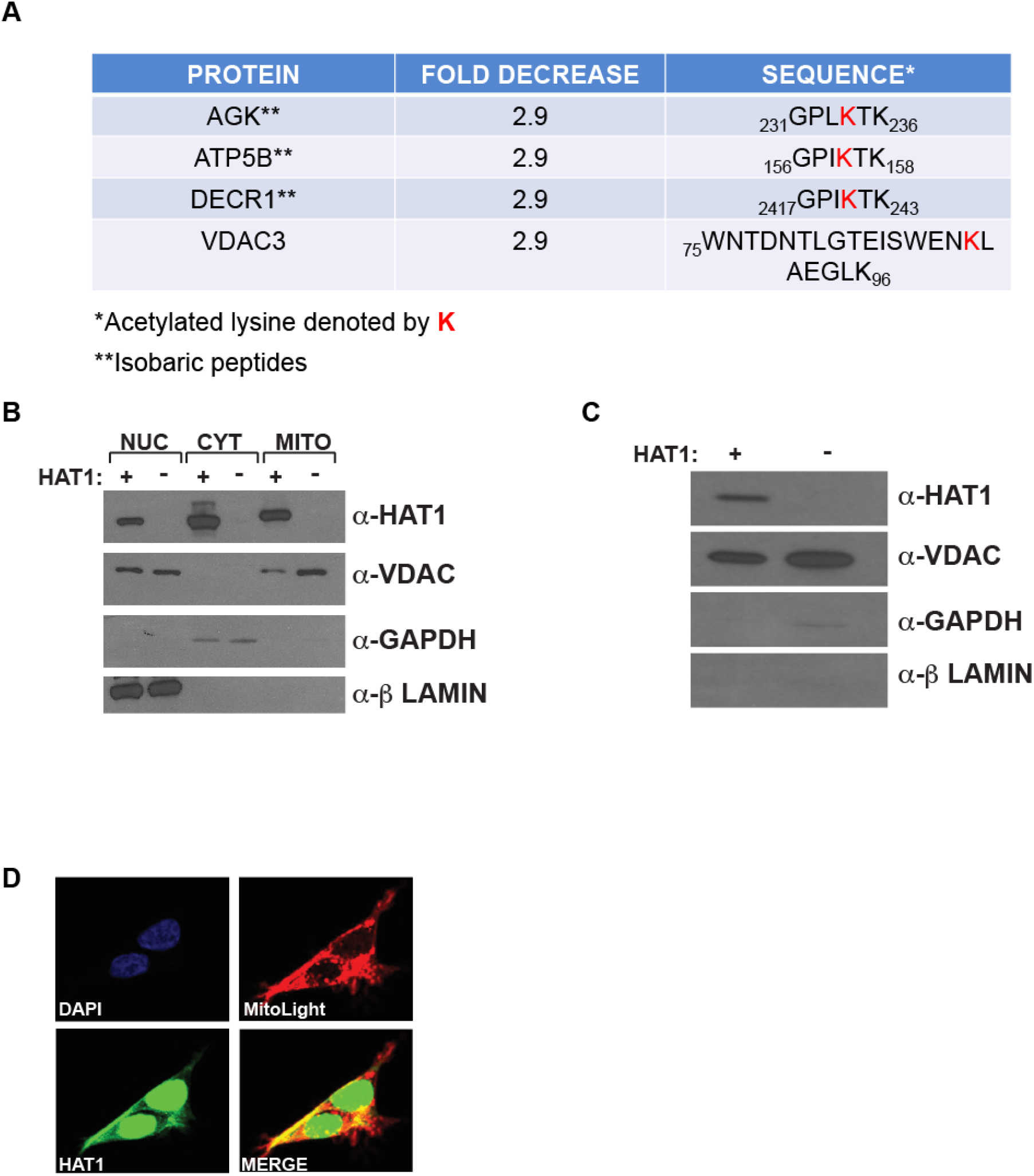
Hat1is required for normal mitochondrial function. A) Subset of mitochondrial proteins whose acetylation decreased at least 2.5 fold in Hat1 KO cells. B) Cells were fractionated int o nuclear, cytoplasmic and mitochondrial extracts. Western blots of the extracts were probed with the indicated markers. VDAC is mitochondrial, GAPDH is cytoplasmic and Lamin is nuclear. C) Mitochondrial were purified by a percoll gradient. Western blot was probed with the indicated antibodies. D) HEK293 cells were transfected with MitoLight expressing vector. Cells were visualized with a confocal microscope and an individual Z-plane is shown.

Even though many mitochondrial proteins are acetylated, there is a debate as to whether mitochondrial protein acetylation results from chemical or enzymatic reactions(48–52). To better understand if it is possible that Hat1 directly acetylates mitochondrial proteins, we examined whether Hat1 localizes in this compartment. We took two biochemical approaches, performing sub-cellular fractionation and mitochondrial purification (Figure 5B). In both cases, Hat1 was detected in the mitochondrial compartment. As expected, the mitochondrial marker VDAC was observed in the mitochondrial fractions but also in the nuclear fraction indicating that there was mitochondrial contamination of the nuclear fraction. However, importantly, nuclear and cytosolic markers, β-Lamin and GAPDH, respectively, were not observed in the mitochondrial fractions suggesting that the Hat1 signal in the mitochondrial fraction is not the result of nuclear or cytosolic contamination. To further confirm this result, we used confocal microscopy to visualize Hat1 and the mitochondrial marker MitoLight. In addition to nuclear and cytoplasmic localization, Hat1 also co-localized with MitoLight in confocal sections (Figure 5C, yellow staining), suggesting that Hat1 may function as a mitochondrial lysine acetyltransferase.

## DISCUSSION

Hat1 was the first lysine acetyltransferase identified but its confirmed substrates are limited to histones. We have used an unbiased proteomics approach to determine how loss of Hat1 influences the mammalian acetylome. This analysis identified 65 proteins whose acetylation is dependent on Hat1. Detailed biochemical experiments will need to be performed to determine whether these proteins are direct substrates of Hat1 or whether Hat1 regulates their acetylation through the regulation of the enzymatic activity of other HATs like CBP. The list of targets that we have identified is likely to be an underestimate of the influence of Hat1 on the mammalian acetylome. In our study, trypsin was used to digest proteins. Trypsin cleaves proteins after lysine or arginine but its activity is blocked by acetyllysine. Hence, loss of Hat1-dependent acetyllysine residues could actually result in an increase in the abundance of some peptides in our study. In addition, trypsin digestion of proteins with high lysine and arginine content can result in peptides that are too small for accurate identification. This is the case with some histones and is likely to be the reason why the validated Hat1 targets, H4 lysine residues 5 and 12, were not detected.

Hat1 was originally isolated from yeast based on its ability to acetylate histone H4 on lysine residues 5 and 12(11). Interestingly, the sequence contexts of H4 lysines 5 and 12 are very similar (GRGK_5_GG and GLGK_12_GG). As the NH_2_-tail of histone H4 is nearly invariant throughout evolution, Hat1 from yeast was capable of acetylating histone H4 from a variety of eukaryotes. Yeast Hat1 was also found to acetylate lysine residue 5 of histone H2A but only from specific species (11,13,33). Examination of the sequence context of histone H2A lysine 5 from a variety of species indicated that Hat1 only acetylated this residue when it was in the context GXG**<u>K</u>**XG, which was hypothesized to be a recognition site for Hat1(11). The subsequent crystal structure of the yeast enzyme demonstrated that there was a structural basis for this recognition site(53). Analysis of the peptides that showed Hat1-dependent acetylation in the current study supports the importance of the glycine residues at positions −3. −1 and +2 but also suggests that these residues are not strictly required. In addition, the Hat1-dependent peptides show a marked preference for an alanine at the +5 position and a glycine at the −7 position. Interestingly, neither of these residues is present at lysines 5 and 12 of H4. However, one or both of the −7 glycine or the +5 alanine are found in canonical mammalian histone H2A and the H2A variants H2A.Z and H2A.J, all of which have decreased acetylation in the absence of Hat1 in our dataset.

A significant fraction of the Hat1-dependent acetylome may be sites that are indirectly regulated by Hat1 through the HATs CBP and Hbo1. Indeed, nearly half of the sites of acetylation that decrease by at least 2.5-fold in Hat1^−/−^ cells were also decreased in CBP^−/−^ cells. There are both direct and indirect mechanisms by which Hat1 may regulate acetylation of the AL of CBP. Hat1 may directly acetylate the lysine residues in the AL of CBP. Alternatively, Hat1 could directly interact with CBP and alter CBP structure or activity to facilitate the autoacetylation of the AL by CBP(34). Hat1 could also interact with other factors that regulate the autoacetylation of the AL, such as RNAs that bind CBP and activate its acetyltransferase activity(54).

Lysine 381 acetylation on p53 (mouse) is both Hat1- and CBP-dependent. Interestingly, Hat1 and p53 share some phenotypes. It was recently shown that mice that are heterozygous for Hat1 (Hat1^+/−^) display premature aging and a significantly shortened lifespan(22). Mice heterozygous for p53 have a similarly shortened lifespan. In addition, heterozygosity of p53 rescues the extreme lifespan decrease of Sirt6^−/−^ mice(55). It will be interesting to determine whether the role of Hat1 in mammalian aging is linked to the acetylation of p53.

While many mitochondrial proteins are acetylated, there is debate as to whether this acetylation is the result of non-specific chemical reactions or is enzyme-catalyzed(48–52). The high levels of acetyl-CoA in the mitochondria certainly suggests that chemical acetylation of mitochondrial is likely to occur. However, there are a number of reports demonstrating the localization of other lysine acetyltransferases to mitochondria(49–51,56,57). Our results, combined with the recent demonstration that Hat1 is required for proper mitochondrial function, raise the possibility that Hat1 may also be involved in mitochondrial protein acetylation(22). A more indirect role for Hat1 in regulating mitochondrial protein acetylation is suggested by a recent report that linked the expression of several genes, including PGC1a, NRF1, NRF and Tfam, to the AMPK-dependent activation of Hat1 in HUVEC cells(29). However, this may be a cell type specific phenomenon, as Hat1 loss in MEFs does not alter the expression of these genes(22). Hat1 doesn’t contain a mitochondrial localization signal, but its translocation to this organelle could occur through the interaction with mitochondrial proteins or mitochondrial transporters. Future studies will focus on determining whether the Hat1-dependent sites of acetylation on mitochondrial proteins are a result of the direct action of Hat1 in the mitochondria. Together the data provided in our study shows that Hat1 influences a wide range of biological process and that its function extends beyond the nuclear compartment.

## ACKNOWLEDGEMENTS

This work was support by a grant form the National Institutes of Health (R01 GM062970).

## AUTHOR CONTRIBUTIONS

P.A.A.G. designed experiments, performed experiments, analyzed data and wrote the manuscript. M.R.P. designed experiments, analyzed data and wrote the manuscript.

